# Cochlear tuning characteristics arise from temporal prediction of natural sounds

**DOI:** 10.1101/2023.10.02.560418

**Authors:** Freddy Trinh, Andrew J King, Ben D B Willmore, Nicol Harper

## Abstract

The cochlea decomposes incoming sound waveforms into different frequency components along the length of its basilar membrane. The receptor hair cells at the apical end of this resonant membrane are tuned to the lowest sound frequencies, with the preferred sound frequency of hair cell tuning increasing near-exponentially along the length of the membrane towards its basal end. This frequency composition of the sound is then transmitted to the brain by the auditory nerve fibers that innervate the inner hair cells. Hair cells respond to a sound impulse with a temporally asymmetric envelope and the sharpness of their tuning changes as the frequency to which they are most sensitive varies with their position along the basilar membrane. We ask if there is a normative explanation for why the cochlea decomposes sounds in this manner. Inspired by findings in the retina, we propose that cochlear tuning properties may be optimized for temporal prediction. This principle states that the sensory features represented by neurons are optimized to predict immediate future input from recent past input. We show that an artificial neural network optimized for temporal prediction of the immediate future of raw waveforms of natural sounds from their recent past produces tuning properties that resemble those observed in the auditory nerve. Specifically, the model captures the temporally asymmetric impulse responses, the tonotopic distribution and variation in tuning sharpness along the cochlea, and the frequency glide polarity of the impulse responses. These characteristics are not captured by a similar model optimized for compression of the sound waveform, rather than prediction. Given its success in accounting for the tuning properties at various processing levels in the auditory and visual systems, this finding for the cochlea provides further evidence that temporal prediction may be a general principle of sensory processing.

## Introduction

Sensory neurons are tuned to diverse but specific features of the environment. While mechanistic models may account for the physical processes behind the tuning properties of sensory neurons, normative models aim to explain what role that tuning plays for the animal and consequently why those properties are seen as opposed to others. Various normative principles have been proposed to explain why particular receptive field properties may evolve; for example, the efficient (Barlow, 1959) or sparse (Olshausen and Field, 1996, 1997; Hateren and Ruderman, 1998; Carlson et al., 2012) use of coding resources for information transmission, or the finding of slowly-varying features (Hyvärinen et al., 2004; Carlin and Elhilali, 2013), given the statistical regularities of natural sensory stimuli. Here, we propose that sensory systems, including the peripheral auditory system, may be governed by a different normative principle, namely temporal prediction (Bialek et al., 2001; Palmer et al., 2015); that is, they are optimized to represent features that can efficiently predict the immediate future of natural sensory input given the recent past input. Optimization for efficient temporal prediction may result in representations that better reflect underlying causes, eliminate irrelevant information, and guide future actions in response to the sensory input (Bialek et al., 2001).

There is evidence for temporal prediction in other sensory systems and at different processing levels. Notably, the temporal prediction principle has been examined in the larval salamander retina (Palmer et al., 2015; Salisbury and Palmer, 2016), where small groups of cells encode near-maximal predictive information. Theoretical work has also found that artificial neural networks optimized for temporal prediction on movies of natural scenes produce model units with receptive fields resembling those of neurons in primary visual cortex (Singer et al., 2018). Furthermore, a hierarchical temporal prediction model, where each subsequent layer is optimized to predict the activity of its input layer, can explain aspects of sensory tuning along the visual pathway, from the retina to the cortex (Singer et al., 2019). Finally, when simple networks are optimized to predict the future of natural sound spectrograms from their past for a dataset of natural sounds, the resulting receptive fields of the hidden units were found to resemble those of neurons recorded in primary auditory cortex (Singer et al., 2018). In this modelling of auditory cortex, the properties of the cochlea were fixed and assumed to be approximated by a spectrogram-like process.

Here, we investigate whether the temporal prediction principle applied to raw natural sound waveforms, rather than spectrograms, can also account for the well-known tuning properties of the cochlea, the site of auditory transduction. The cochlea acts as a frequency analyser and consists of a coiled tube divided lengthwise into three fluid-filled chambers (reviewed in Fettiplace, 2017). The central chamber is bounded on one side by the basilar membrane, which supports the Organ of Corti, where the receptor hair cells are located. The mechanical properties (notably stiffness) of the basilar membrane vary smoothly along the length of the cochlea, resulting in a gradient of frequency tuning. The sound frequency to which each region of the basilar membrane is tuned increases approximately exponentially from apex to base (Liberman, 1982), so that relatively more length is dedicated to low frequencies than to high frequencies. Vibration at a point on the basilar membrane is detected by the hair cells in the corresponding part of the Organ of Corti, and then transmitted to the brainstem by the auditory nerve fibers.

The impulse response of a point on the basilar membrane is largely inherited by the corresponding inner hair cell and auditory nerve fibers, such that auditory nerve fiber activity can be used to assess cochlear tuning (Narayan et al., 1998). In this paper, we focus on the impulse response function inherited from the cochlea, as measured from the firing of auditory nerve fibers. We will refer to these inherited impulse responses as ‘cochlear filters’. This tuning can be approximated by a gammatone filter (de Boer, 1975; Carney, 1993), which (in the time domain) is a sinusoidal function characterized by a temporally asymmetric envelope with a fast onset and a more gradual offset (Goblick and Pfeiffer, 1969). The absolute filter bandwidth increases with the characteristic frequency of the auditory nerve fibers, i.e. the sound frequency where the minimum response threshold is found (Glasberg and Moore, 1990; Walker et al., 2019). However, the relative bandwidth decreases with characteristic frequency, that is sharpness of tuning expressed as the quality factor (Q = characteristic frequency/bandwidth) increases (Evans, 1972; Oxenham and Shera, 2003; Joris et al., 2011; Sumner and Palmer, 2012).

The representation of sound in the cochlea has previously been investigated by models optimized for efficient coding of sound (Lewicki, 2002; Smith and Lewicki, 2006). These models produced units with impulse responses that were similar to cochlear filters in many respects, showing sinusoidal ringing within a temporally limited envelope, and a range of frequency tuning across units. However, the envelopes of the impulse responses in Lewicki (2002) were temporally symmetrical in form, in contrast to the asymmetric temporal envelopes of cochlear filters. The model of Smith and Lewicki (2006) produced temporally asymmetric gammatone-like impulse responses, but the inference process used by the model did not respect causality, as it employed an iterative process that required all time points comprising a long waveform to be available simultaneously.

We asked whether the temporal prediction principle could capture features of cochlear tuning. Our initial approach made relatively few assumptions about the components of the cochlea. We modelled the cochlea as a bank of filters that were optimized to represent the features that best predicted the immediate future of natural sound waveforms from their recent past. We found that this simple model could reproduce many of the tuning properties of cochlear filters.

## Results

### The cochlear temporal prediction model

To examine whether optimization for temporal prediction could explain cochlear tuning properties, we trained a simple neural network model to predict the immediate future of sound waveforms from their recent past. The model was a feedforward neural network with one layer of hidden units (Fig. 1) (see Methods for more details). The input layer consisted of 128 units fully connected to 64 hidden units, and the hidden units were fully connected to 16 output units. The input was the immediate past waveform, and the activity of each hidden unit was a weighted sum of the input followed by a nonlinear transform using the scaled hyperbolic tangent function. The output weights of the network produce the output unit activity, which is a linear estimate of the upcoming future from the activity of the hidden units. We assumed that the code used by the cochlea can be approximately linearly decoded to provide a prediction of the future input. This is consistent with the observation that decoding stimuli from neural responses is often performed surprisingly well by a linear transformation (Eliasmith and Anderson, 2002).

**Figure 1.**
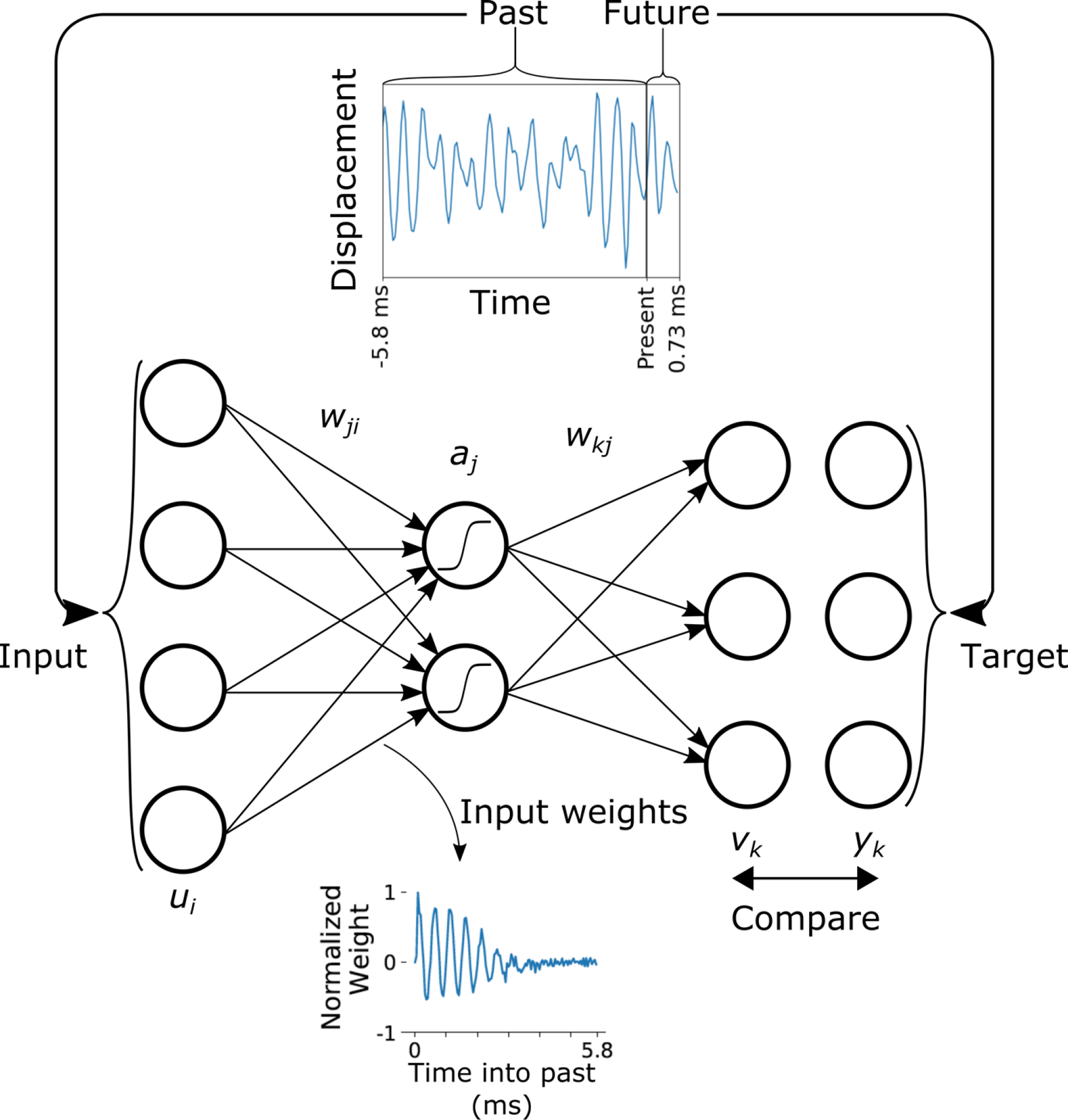
Schematic of the temporal prediction model of the cochlea. A large dataset of natural sound waveforms (sampled at 22,050 Hz) was divided into snippets of 144 samples (6.5 ms) and presented to the network in random order. The first 128 samples (5.8 ms) of each snippet were provided as inputs to a feedforward neural network, and the network was optimized to perform temporal prediction by predicting the 16 future samples (0.73 ms) following the input samples. Each input unit, *u*_*i*_, represents the waveform displacement at a particular input sample with index *i*, while each output unit, *v*_*k*_, predicts the displacement at a future sample, with index *k*. The nonlinear hidden units are indexed by *j* and have activity *a*_*j*_. The linear weights, *w*_*ji*_ and *w*_*kj*_, and the biases, *b*_*j*_, and *b*_*k*_, were optimized to produce a prediction, *v*_*k*_, of the future, *y*_*k*_, given the past, *u*_*i*_, over the dataset of natural sounds. The upper inset figure shows an example sound snippet representing the past, *u*_*i*_, and the future, *y*_*k*_, which served as input and target, respectively, for the temporal prediction objective. An example of the tuning properties learned by the model is shown in the lower inset figure. The incoming weights, *w*_*ji*_, to hidden unit *j* can be interpreted as the impulse response of this hidden unit. Note for clarity we have dropped the snippet index *n* that we use in the Methods from *u*_*i*_, *a*_*j*_, *v*_*k*_ and *y*_*k*_.

To train the model to predict future input, we used a set of sound recordings representative of the natural auditory environment. This dataset included birdsong, animal vocalizations, environmental sounds, human speech by children and adults, and baby vocalizations. The stimuli were linearly filtered to mimic the transformation performed by the middle ear (see Methods). We then added Gaussian noise to the input (9 dB signal-to-noise ratio, SNR), which has been found to aid in the capturing of sensory tuning properties by temporal prediction models (Singer et al., 2018). The model parameters were trained to predict the waveform over a brief timespan into the future (16 samples = 0.73 ms) from the present, given the recent past (128 samples = 5.8 ms) of the waveform immediately before the present. The model was trained using backpropagation to minimize the mean squared error (MSE) between the future waveform and the prediction of the waveform by the model. Once the model was trained, we interpreted the pattern of input weights to each hidden unit as being analogous to the impulse response of a specific site along the basilar membrane.

### Properties of the model when trained on natural sounds

The trained temporal prediction model produced substantial impulse responses for most of the hidden units (51 out of 64) (Fig. 2). Hidden units with impulse response power below 1% of the average impulse response power were deemed not substantial and removed from further analysis. The impulse responses showed the characteristic sinusoidal ringing elicited by the basilar membrane with a rapid onset followed by a more prolonged offset. The envelopes were temporally asymmetric, with most energy at recent time lags and the energy decaying into the past. Hence, the model impulse responses replicate characteristics of cochlear filters. However, the onset of the sinusoidal ringing was generally faster for the model impulse responses than in real cochlear filters. Unlike biological impulse responses, some model impulse responses also showed a strong preference for the most recent stimulus sample, as seen by a click-like rise. Figure 3 illustrates the similarity between representative impulse responses of the trained model (Fig. 3A) and impulse responses recorded from auditory nerve fibers of anesthetized cats (dataset from Carney et al., 1999) (Fig. 3B).

**Figure 2.**
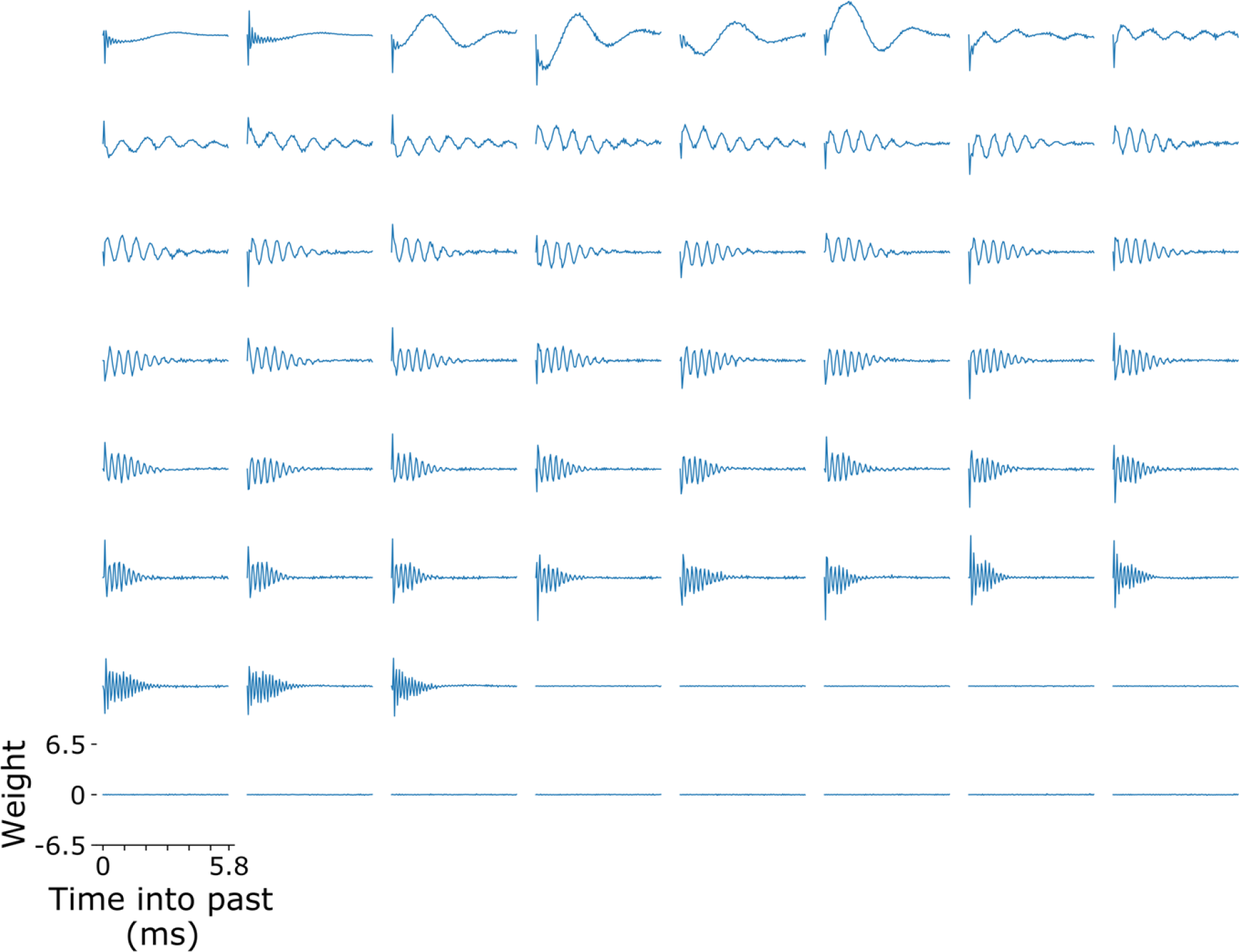
All impulse responses for the temporal prediction model. The impulse responses of all 64 hidden units of the model sorted by center frequency, which was determined by the position of its peak value in frequency space. The impulse response for hidden unit *j* is the weight, *w*_*ji*_, from all input units, *i*, with each input unit providing the input from a particular time point in the past. If the power of an impulse response failed to exceed a threshold of 1% of the average model impulse response power, it was removed from further analysis (last 13 hidden units shown).

**Figure 3.**
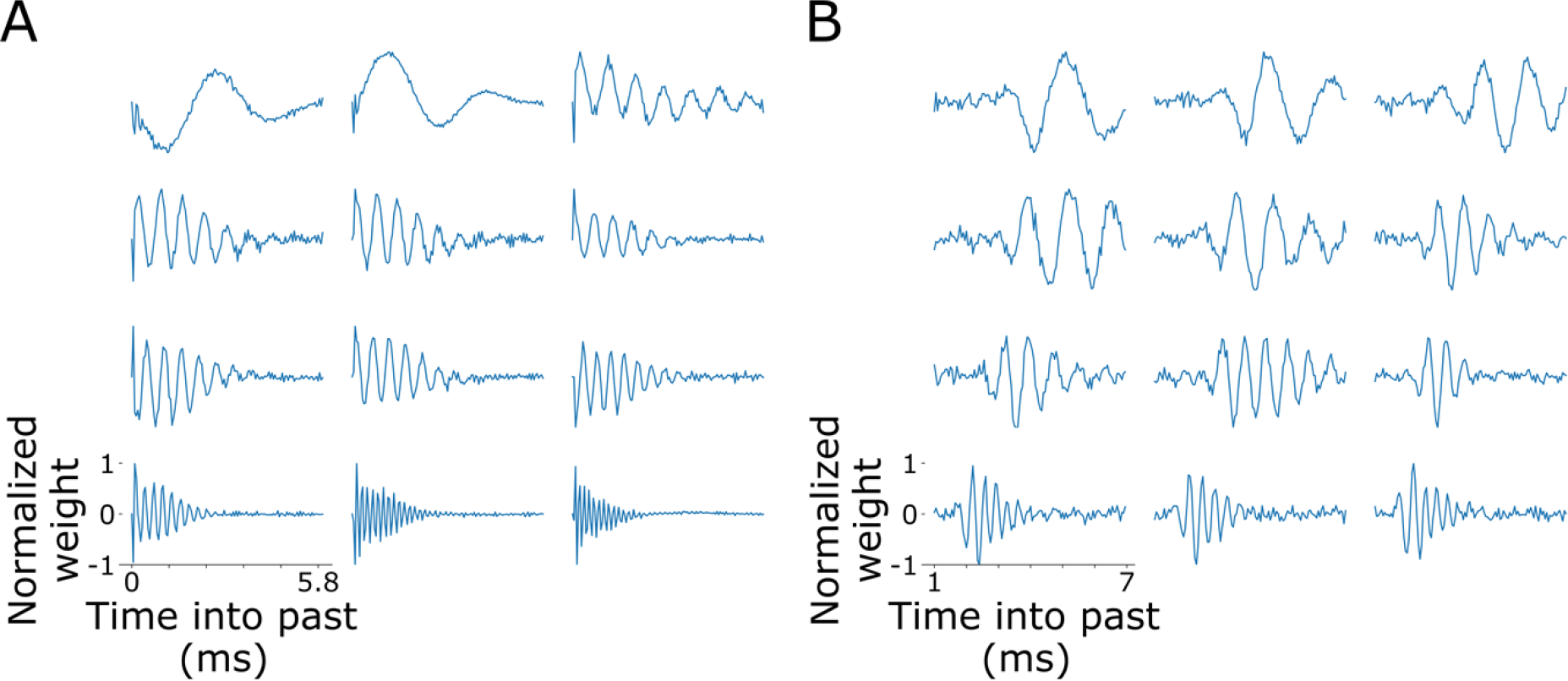
Impulse responses of representative hidden units of the temporal prediction model and auditory nerve fibers. Twelve examples are shown in each case, illustrating the similarity between them. (A) Impulse responses of the network model. The impulse response for each hidden unit was normalized to its respective maximum absolute value. (B) Impulse responses of auditory nerve fibers recorded in anesthetized cats and obtained using reverse correlation (Carney et al., 1999).

We determined the center frequency of each hidden unit as the peak of its impulse response in frequency space. As our model units only had a simple compressive nonlinearity, their center frequency will be the same as their best frequency at each sound intensity and their characteristic frequency. Each hidden unit displayed sharp frequency tuning, and when the units were ranked by their center frequency, they smoothly spanned a range from ∼175 Hz up to ∼7000 Hz (Fig. 4A). We used an exponential equation that describes the tonotopic cochlear map in mammals (Greenwood, 1961, 1990, 1996; Robles and Ruggero, 2001) (Fig. 4B) to fit the center frequency (*f*) as a function of the units’ rank order (Fig. 4B). This is given by *f* = *A*(*10*^*Cz*^ *+ B*), where *z* is the distance from the cochlear apex scaled from 0 to 1, and *A, B* and *C* are the fitted parameters. Our model does not provide an absolute measure of distance from the cochlear apex, but we took each unit, ordered by center frequency, to represent an equal-sized adjacent space along the basilar membrane. Thus, the units represent a contiguous section somewhere along the basilar membrane. We took z = 0 at the position of the lowest-frequency model unit (not the apex), and z = 1 to be the position of the highest-frequency model unit (not the base), with each unit in between assigned a value of z proportional to its rank order r between these limits, that is *z* = (*r*-1)/(*r*_max_-1), where *r*_max_ = 51 is the number of units with substantial impulse responses. The exponential function provided a better fit than a linear fit (F test, *F*(49,48) = 320.05, *p* = 1.11×10^-16^). Hence, the range of impulse responses in the model approximately captured the smooth, near-exponential arrangement of tuning frequency exhibited by the cochlea (Greenwood, 1961, 1990, 1996; Liberman, 1982; Robles and Ruggero, 2001).

**Figure 4.**
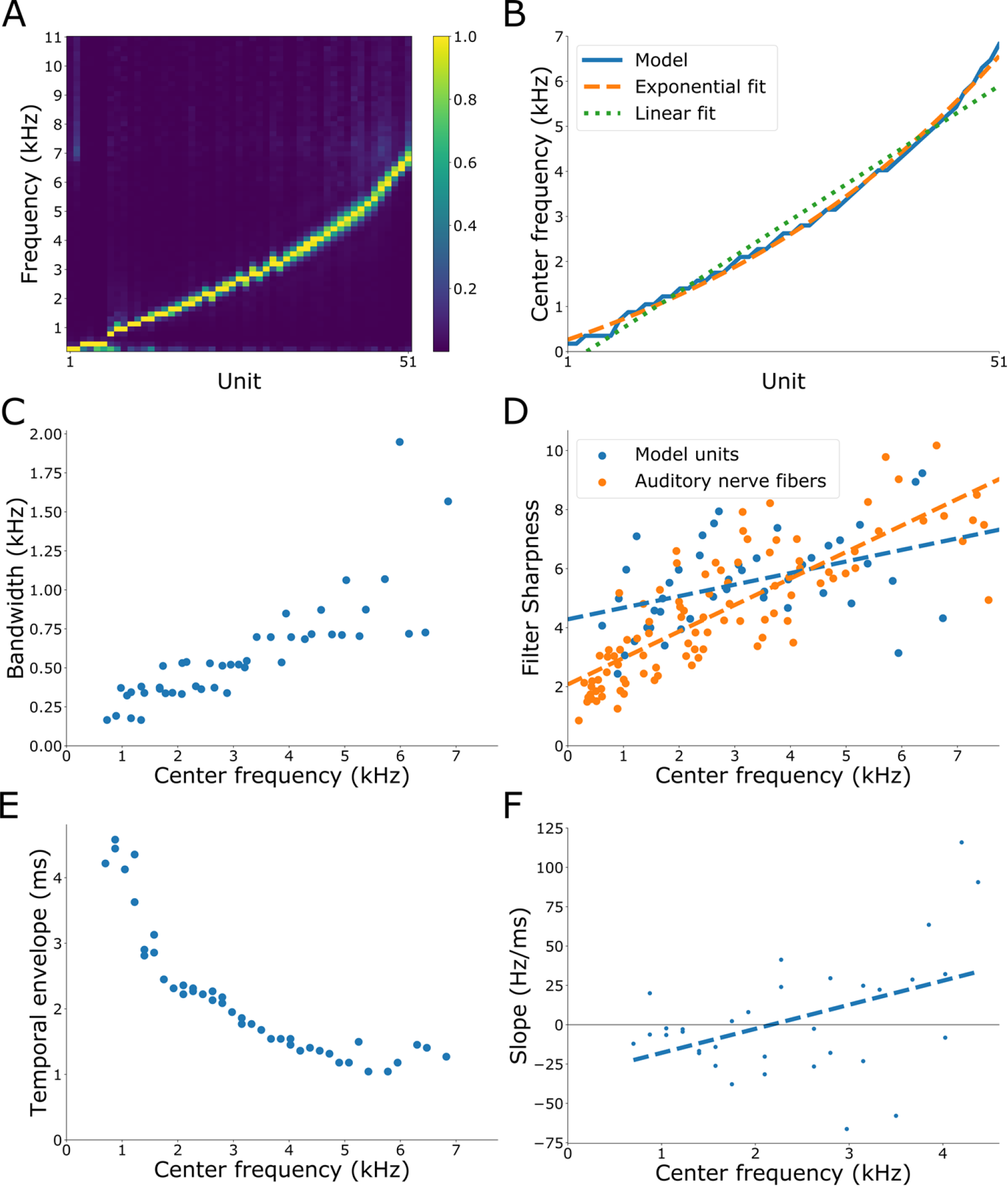
Impulse response characteristics of the temporal prediction model. (A) The normalized power spectrum of the impulse responses of each hidden unit of the feedforward model, with the units ranked by center frequency. The color-bar represents the normalized power. (B) The center frequency of each model unit as a function of hidden unit rank. The hidden units are interpreted to span a contiguous section of the basilar membrane in tonotopic order, with each unit evenly spaced along this section. The model centre frequencies are fitted with an exponential equation that has been used to describe the relationship between distance along the basilar membrane and characteristic frequency (Greenwood, 1961, 1990, 1996; Robles and Ruggero, 2001). The best linear fit of the model center frequencies is also shown. (C) The impulse response bandwidth of each hidden unit plotted against center frequency. The impulse response bandwidth was estimated at 10 dB below the spectral peak of the impulse response. (D) The filter sharpness (Q_10_) of each hidden unit plotted against center frequency. The filter sharpness was determined by dividing the center frequency by the impulse response bandwidth. Blue dots indicate data points taken from the temporal prediction model. Orange dots indicate data points recorded from cat auditory nerve fibers; the data were extracted from Figure 2 of Evans (1975). The best linear fit of both the model and experimental data are shown as dashed lines in their respective color. (E) The temporal envelope of each hidden unit plotted against center frequency. The temporal envelope of each impulse response was determined by the shortest time window possible that captured 95% of the power of the impulse response. (F) For each impulse response, the instantaneous frequency was measured at different points in time using the zero-crossing method, and then fitted with a straight line to obtain the slope of its frequency glide (Carney et al., 1999). See Methods for details. Each point is the slope of the frequency glide of an impulse response, plotted against the center frequency of that impulse response, for the range of neuronal best frequencies measured in Carney et al. (1999). Negative slope values indicate a decrease in frequency as the impulse response progresses (a downwards glide); positive slope values indicate an increase (an upwards glide). The dashed line is the best linear fit.

The bandwidth of the impulse responses increased as the center frequency increased (Pearson correlation coefficient = 0.816, *p* = 7.01x10^-12^), as seen for cochlear filters (Fig. 4C) (Moore, 1986; Glasberg and Moore, 1990). The relative sharpness of each impulse response was estimated by dividing the center frequency of the impulse response by its bandwidth at 10 dB below the spectral peak of the impulse response, thereby generating Q_10_ values that could be compared to physiological data. In the auditory nerve, the Q_10_ increases with characteristic frequency (Pearson correlation coefficient = 0.816, *p* = 2.43x10^-24^ for the cat data in Evans, 1975 (Fig. 4D). The filter sharpness of the hidden units of our model similarly increased with centre frequency (Pearson correlation coefficient = 0.44, *p* = 0.00248), which matched the experimental findings fairly well over the frequency range of the hidden units (Fig. 4D). The joint distribution of the model hidden units’ center frequencies and sharpness measures was not significantly different from that of the physiological data (Wald-Wolfowitz test, W = 0.393, *p* = 0.6527) (Friedman and Rafsky, 1979; Monaco, 2014). The temporal envelope of an impulse response is the temporal span that captures 95% of the impulse response’s power (Lewicki, 2002). We found that this decreased for the model hidden units with increasing center frequency (Pearson correlation coefficient = -0.86, *p* = 7.592x10^-14^) (Fig. 4E).

Finally, we investigated the instantaneous tuning properties of the model impulse responses. Detailed investigation of the impulse responses of auditory nerve fibers of the cat suggests that the frequency tuning of the impulse response changes slightly over its duration; rather than being exactly a gammatone, the impulse response instead often exhibits a glide either up or down in frequency (Carney et al., 1999). This time-dependent frequency tuning (‘instantaneous frequency’ as a function of time) can be measured by looking at the timing of the zero-crossings of the impulse responses over time (See Methods and Supplemental Fig. 1). Carney et al. (1999) found that auditory nerve fibers tuned to low frequencies (<1500 Hz) tend to decrease or remain steady in instantaneous frequency over time (downward frequency glides), whereas those tuned to higher frequencies (>1500 Hz) tend to increase in instantaneous frequency over time (upwards frequency glides). Thus, the slope of instantaneous frequency over time for impulse responses goes from negative to positive as the preferred sound frequency of auditory nerve fibers increases. We plotted these slopes as a function of center frequency for the impulse responses of the temporal prediction model (Fig. 4F). This was done over the same range of auditory nerve fiber best frequencies measured by Carney et al. (1999) to facilitate comparison with the experimental data and because estimates of instantaneous frequency by the zero-crossing method are inaccurate above half the Nyquist frequency (Hoeks et al., 1984). Consistent with the cat auditory nerve recordings findings (Carney et al., 1999), we found that the slopes of the model units increased with their center frequency (Pearson correlation coefficient 0.44, *p* = 0.0086). Thus, although the magnitude of the slopes was typically several times smaller than that seen in the experimental data, the same general trend of slope polarity going from negative to positive with increasing center frequency was seen in the temporal prediction model.

### The role of noise

We found that the addition of weak Gaussian noise to the input was necessary for the model to learn cochlea-like impulse responses (Fig. 3). An absence of noise in the training set produced a representation that differed from that found in the cochlea and auditory nerve fibres (Fig. 5A,B). The impulse responses did not reproduce gammatone impulse responses, and did not exhibit a gradual offset. Instead, the impulse responses displayed high-frequency oscillations with sequences of staggered onsets and offsets, and some contained an additional underlying low frequency oscillation (Fig. 5A). Most impulse responses were not sharply tuned in frequency space and the center frequencies of the impulse responses were separated onto opposite ends of the frequency space with virtually no tuning to the mid-range frequencies (Fig. 5B).

**Figure 5.**
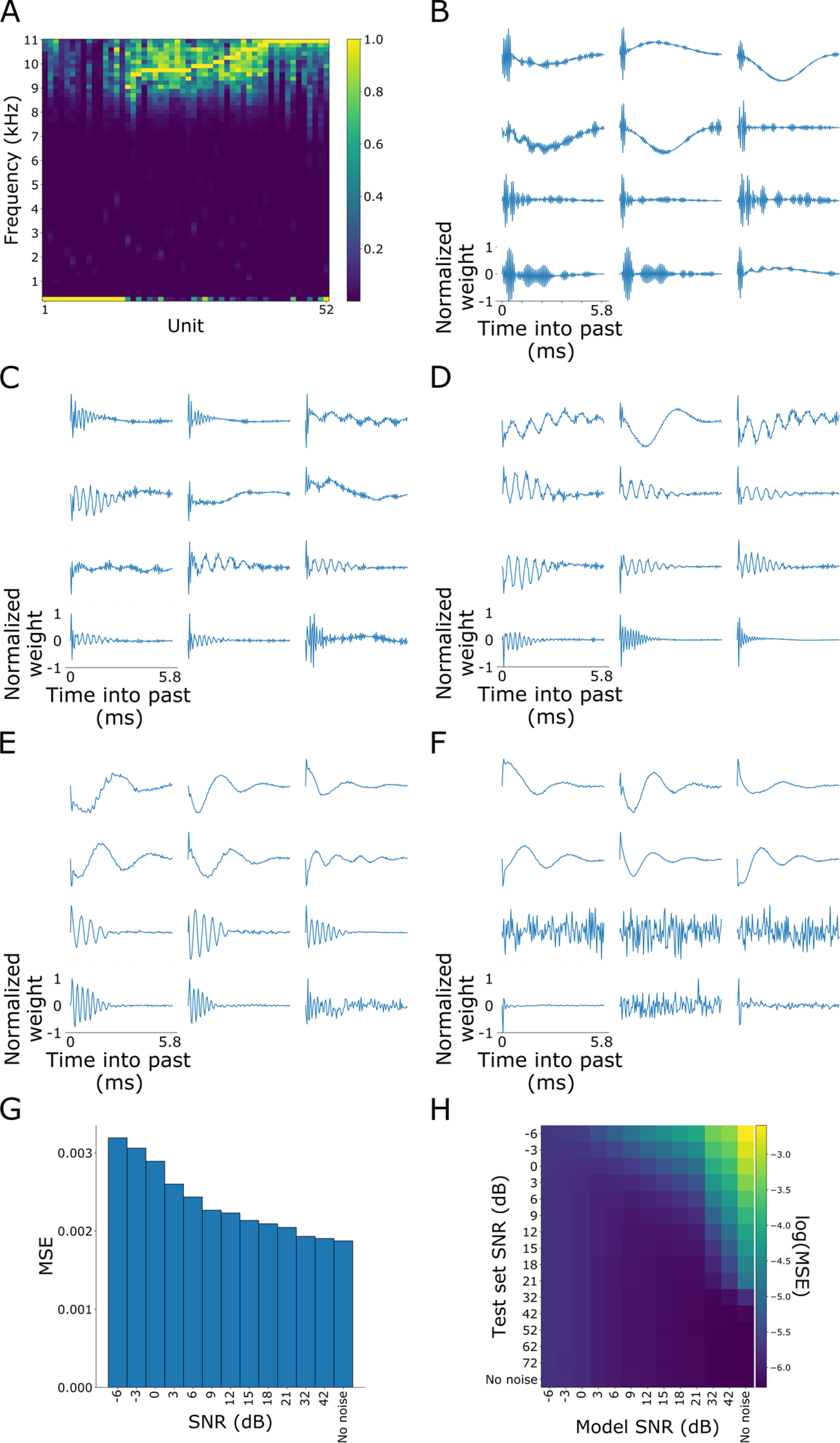
Impulse responses of temporal prediction models trained with various levels of noise added to the input. **(A**,**B)** Impulse response characteristics of the temporal prediction model trained on noiseless sound. **(A)** The normalized power spectrum of the impulse response of the hidden units of the feedforward model. The color-bar represents the normalized power. **(B)** Example impulse responses of the network model trained on noiseless sound. The impulse response for each hidden unit was normalized to its respective maximum absolute value. **(C-F)** Impulse responses of models trained on a signal-to-noise ratio (SNR) of 21, 15, 3 and 0 dB, respectively. The impulse response for each hidden unit was normalized to its respective maximum absolute value. **(G)** Prediction performance (MSE, mean squared error) of temporal prediction models trained with various levels of noise added to the input (but not the target). In each case, the models were tested on a held-out test set consisting of natural sounds without added noise. **(H)** As G, but models were tested using inputs consisting of natural sounds with various levels of noise added. The bottom row (noiseless test data) is the same as G.

To determine the effect of input noise on the prediction performance of our network, we explored the effect of applying noise to the input at a range of different signal-to-noise ratios (SNRs) (Fig. 5C-F). Across a range from 3 to 15 dB SNR, hidden units exhibited cochlea-like tuning (Fig. 5D, E). The similarity between the hidden units and biological tuning deteriorated for noise levels outside this SNR range (Fig. 5C, F). We speculated that the noise might act as a regularizer, in which case we might expect that the added noise should enable the temporal prediction model to better predict future input when evaluated on a held-out noiseless test set. In fact, we found that the no-noise case provided the best predictions on the noiseless test set, suggesting that the input noise does not regularize the model (Fig. 5G). However, when we added noise to the test set inputs, we found that models trained in the range 3-15 dB SNR were particularly good at predicting the future across a range of test SNR levels (3-72 dB and no noise) (Fig. 5H). This suggests that the cochlea may use a representation that is robust to the presence of varying levels of internal and external noise.

We also examined whether where in the model the noise was added was important for getting gammatone-like tuning curves. When Gaussian noise was added to the future targets rather than the past input, there was a deterioration of the biological similarity of the model (Supplemental Fig. 2A-B), similar to what was seen in the noiseless case. However, adding noise on both input and target resulted in a similar representation to the model trained on noisy input alone (Supplemental Fig. 2C-D), though there were fewer active units (34/64 hidden units were active) and a subset of these lacked sharp frequency tuning. We conclude that noise added to the input was the crucial factor in reproducing cochlear tuning properties.

To investigate the possibility that the noise enabled avoidance of local minima and saddle points during gradient descent, we took the network trained with 9 dB SNR and then trained it further with the noiseless stimuli. However, the resulting network showed the same results as the network that was simply trained with the noiseless stimuli (Supplemental Fig. 2E-F). Finally, we asked whether the reason why noise was necessary to obtain cochlea-like model features was due to it acting as a form of data augmentation. We trained a network on the same dataset without any added noise, but with approximately 10 times more sound snippets. We did this by choosing 6.5 ms snippets from the sound clips with onsets that were spaced by 0.65 ms, rather than 6.5 ms, providing many more snippets from the same sound clips, which now overlapped. This model still produced impulse responses that were not gammatone-like and without the characteristic cochlear tonotopic mapping (Supplemental Fig 2G-H). This suggests that the addition of noise did not function as data augmentation in order to settle on cochlea-like properties.

### The cochlear autoencoder model

We next also asked whether temporal prediction itself was crucial for our model to learn cochlea-like tuning. It is possible that a network that merely compresses auditory stimuli is sufficient to produce cochlea-like tuning, and that temporal prediction *per se* is not required. Two previous models based on the principles of efficient coding have had some success in reproducing aspects of cochlear filters (Lewicki, 2002; Smith and Lewicki, 2006), though they have some limitations as outlined in the Introduction. Efficient coding seeks to use coding resources efficiently to represent all the incoming sensory input, and can be viewed as a form of compression of the input (Chalk et al., 2018). This is in contrast to temporal prediction, where incoming sensory input is only represented to the degree that it is predictive of future input. To investigate the extent to which shared characteristics between the temporal prediction model and cochlear features can be explained by compression rather than prediction, we trained an autoencoder neural network (Supplemental Fig. 3). The autoencoder model is identical to the temporal prediction model (Figs. 1-4), apart from the output layer, which recreates the input rather than predicting the immediate future.

Unlike the temporal prediction model, the autoencoder was unable to capture the same range in frequency tuning and did not display sharp frequency tuning apart from a small subset of hidden units (Fig. 6A). The span of the center frequencies of the hidden units did not capture the exponential relationship between center frequency and basilar membrane position seen in the cochlea, and was best fitted linearly (F test, *F*(62,61) = 0.34, *p* > 0.99) (Fig. 6B). A few somewhat gammatone-like impulse responses were produced, but without the temporal asymmetry exhibited by biological cochlear filters or the temporal prediction model (Fig. 6C). Furthermore, the bandwidth of the impulse responses did not uniformly increase as a function of center frequency (ρ = 0.19, *p* = 0.13), (Fig. 6D). The filter sharpness increased as a function of center frequency (ρ = 0.6, *p* = 1.5×10^-7^), but overshot the biological data across the frequency range (Wald-Wolfowitz test, W = -1.34, *p* = 0.0901) (Fig. 6E). Finally, in contrast to cochlear filters, the temporal envelope of the autoencoder impulse responses was not correlated with the center frequency (Pearson correlation coefficient = -0.044, *p* = 0.731) (Fig. 6F).

**Figure 6.**
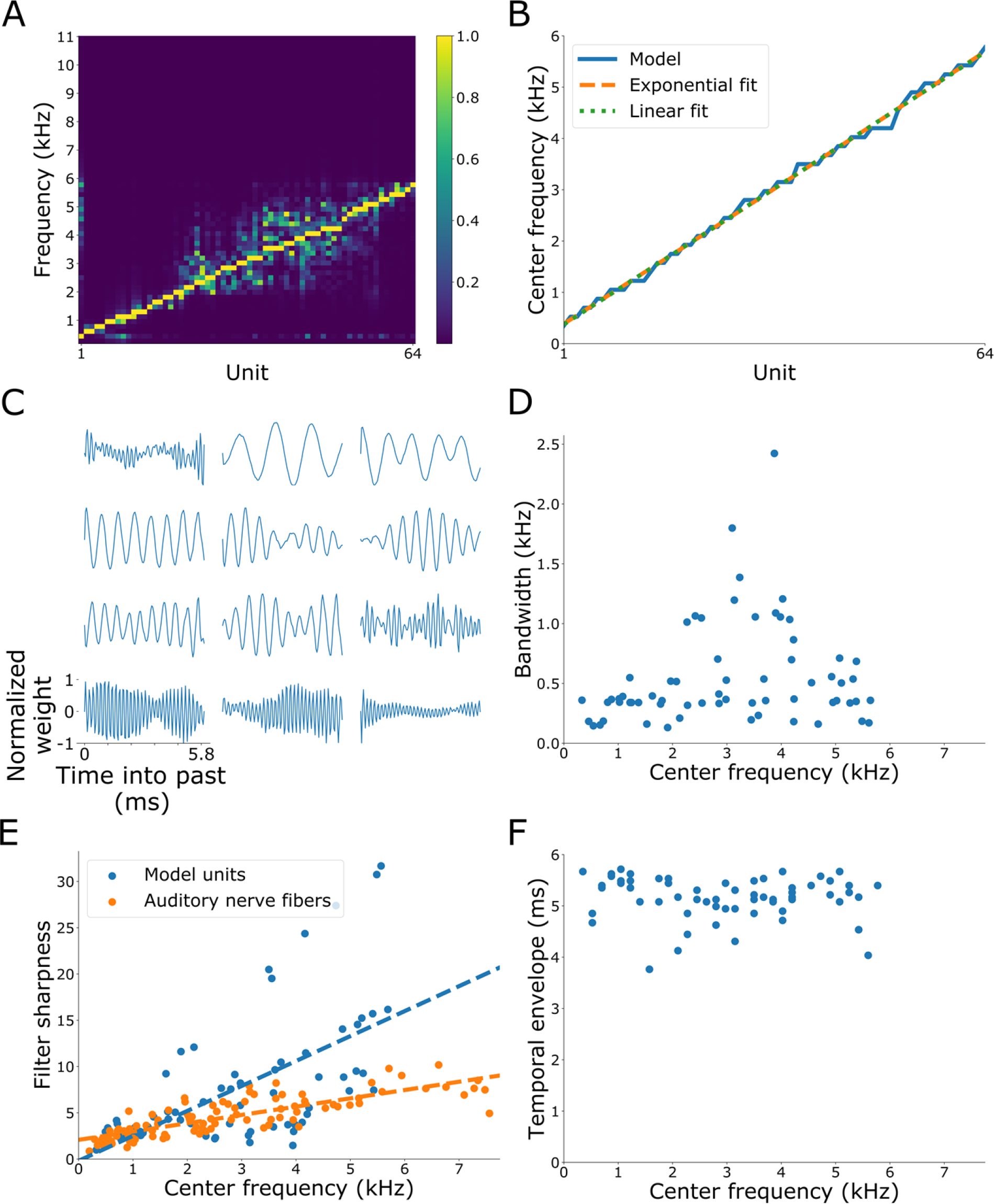
Impulse response characteristics of the autoencoder model. **(A)** The normalized power spectrum of the impulse response of the hidden units of the feedforward model. The color bar represents the normalized power. **(B)** The center frequency of each model unit as a function of hidden unit rank. The hidden units are interpreted to span a contiguous section of the basilar membrane in tonotopic order, with each unit evenly spaced along this section. The model center frequencies are fitted with a logarithmic equation that has been used to describe the relationship between characteristic frequency and distance along the basilar membrane (Greenwood, 1961, 1990, 1996; Robles and Ruggero, 2001). The best linear fit of the model center frequencies is also shown. **(C)** Impulse responses of the network model. The impulse response for each hidden unit was normalized to its respective maximum absolute value. **(D)** The impulse response bandwidth of each hidden unit plotted against center frequency. The impulse response bandwidth was estimated as a 10 dB decrease on either side of the spectral peak of the impulse response. **(E)** The filter sharpness of each hidden unit plotted against center frequency. The filter sharpness was determined by dividing center frequency by impulse response bandwidth at 10 dB below the peak response (Q_10_). Blue dots indicate data points taken from the temporal prediction model. Orange dots indicate data points taken from cat auditory nerve fibers (Evans, 1975). **(F)** The temporal envelope of each hidden unit plotted against center frequency. The temporal envelope of each impulse response was determined by the shortest time window possible that captured 95% of the power of the impulse response.

The autoencoder was trained on the 9 dB SNR sounds. For completeness we also trained the autoencoder on the noiseless sounds. The results were similar to the autoencoder trained on the noisy sounds, but the impulse responses were even less tightly frequency tuned (Supplemental Fig. 4).

## Discussion

We asked whether temporal prediction applied to natural sounds might be a computational principle underlying stimulus representation in the cochlea. To investigate this, we optimized a feedforward network with one hidden layer to predict the immediate future of sound waveforms given their recent past. The model was able to capture several features of cochlear tuning. The hidden units showed a wide range of frequency tuning, with more units tuned to lower frequencies than higher frequencies, consistent with the near-exponential dependence of characteristic frequency on basilar membrane position (Greenwood, 1961, 1990, 1996; Liberman, 1982). The impulse responses of the hidden units had gammatone-like temporal profiles and resembled those recorded from cat auditory nerve fibers (Fig. 2 and 3). More specifically, the impulse responses were temporally asymmetric with a rapid onset followed by a slower offset. Consistent with the biology, the absolute bandwidth and filter sharpness (expressed as Q_10_) of the model impulse responses increased as a function of centre frequency, while the temporal envelope decreased (Fig 4) (Evans, 1975; Moore, 1986; Lewicki, 2002). Additionally, the model exhibited a dependence of the polarity of the frequency glides of the impulse responses on centre frequency, as seen in the real auditory nerve (Carney et al., 1999). In contrast to the temporal prediction model, we found that a simple autoencoder failed to reproduce many of these cochlear features. Together, these findings suggest that optimization for temporal prediction on natural sound waveforms can account for many aspects of cochlear sound processing.

### Factors important for the model

The addition of Gaussian noise to the input at ∼3-15 dB SNR proved to be needed for the model to replicate cochlear features (Fig 5 and 6). We investigated three technical reasons that might explain why the noise was needed: regularization, data augmentation, and avoiding local minima or saddle points. The addition of noise to network input can act as regularization that reduces overfitting and helps with generalization of a model (Sietsma and Dow, 1991). It has been shown that, under certain assumptions, adding noise is equivalent to Tikhonov regularization (a generalization of L2 regularization) when the noise amplitude is small (Bishop, 1995, 2006). If this is the reason why noise was beneficial in our model, we would expect noise to improve prediction performance on a held-out noiseless dataset. This was not the case (Fig. 5G), however, suggesting that noise is not acting as a regularizer.

Another possibility is that the addition of noise approximates having a larger training set; addition of noise to a training set is often used as a type of data augmentation in machine learning (Goodfellow et al., 2016). If this were the case in our model, we might expect that a larger, noiseless data set would produce similar results to those found for the dataset with noise added. This was not the case (Supplementary Figure 2E-F). Finally, noise can help during gradient descent to avoid local minima and saddle points. To test whether this was the case in our model, we took the model trained with noise and then trained it further on the same dataset but without noise. If the noiseless model was finding a poor local minimum or saddle point, then pre-training with noisy data should avoid this. However, even with pre-training (which produced gammatone-like impulse responses), subsequent training on noise-free data resulted in a model with non-gammatone impulse responses. Thus, while our exploration of this question is not exhaustive, these results suggest that the noise does not seem to be simply enabling better training of the network.

This leaves open the possibility that the noise we added mimics some form of noise that is unavoidably present in the middle or inner ear or in the environment. Thermal noise may induce stochasticity at the middle ear (Harrison, 2009), basilar membrane (Nuttall et al., 1997) and hair cell stereocilia (Kozlov et al., 2012). Furthermore, noise from cardiovascular and respiratory muscle activity (Ren et al., 1995; Nuttall et al., 1997) and other muscle activity, such as chewing (Bárány, 1938; Huxley, 1990), may also have an impact at low sound frequencies (Nuttall et al., 1997; Kirk and Smith, 2003). There is also substantial stochasticity in the vesicle release process of the inner hair cell ribbon synapse (Heil and Peterson, 2017). Given the relative quietness of many natural environments (Kirk and Smith, 2003), this internally-generated noise may be of relevance. However, the presence of externally-generated nuisance noise, including wind (Chung, 2012), reverberation (Traer et al. 2016) and other background natural sound textures, such a rain or flowing water (McDermott et al. 2011; Mishra et al. 2021), may also have influenced the evolution of the auditory system. Hence, it may be that the cochlea is optimized for temporal prediction of natural sounds in moderately noisy backgrounds with noisy biological mechanisms. This is supported by the fact that training in the 9-15d B SNR range appears to allow reasonably good prediction of the test set over a wide range of SNRs (from no noise to 3 dB SNR) (Fig. 5H).

Our results also suggest that compression of the sound waveform is not sufficient to account for cochlear tuning. An autoencoding, compressive network trained to recreate its natural sound waveform input did not emulate cochlear features well, as the autoencoder did not produce tightly-tuned, appropriately asymmetric, and exponentially-distributed impulse responses. This suggests that the cochlear features produced by temporal prediction are not solely due to compression and are a consequence of encoding the predictive features of the natural soundscape.

### Comparison to other models

Models of sensory coding have previously been based on several other normative principles, one being efficient coding (Barlow, 1959). Efficient coding postulates that sensory systems are optimized to represent natural stimuli by minimizing redundant activity between transmitting neurons (Lewicki, 2002). Cochlear models optimized to efficiently represent natural sound waveforms have been shown to produce filters that resemble the impulse response of auditory nerve fibres (Lewicki, 2002; Smith and Lewicki, 2006). However, the efficient coding approach used by Lewicki (2002) did not capture the asymmetry of biological impulse responses. The encoding algorithm of Smith and Lewicki (2006) did produce asymmetric impulse responses, but is also limited in its biological plausibility by producing temporally-sparse spikes, each with a scalar value, rather than binary spikes (as in the auditory nerve) or a continuous real number (as with the inner hair cell membrane potential). Also, while this encoding algorithm produces spikes sequentially over time, it treated time as a dimension viewed all at once, with the timing of past spikes being able to depend on those of future spikes, something inconsistent with biology. Our model has addressed some of these shortcomings by using temporal prediction as a governing principle. The impulse responses from the temporal prediction model capture the temporal asymmetry of cochlear filters, and its encoding method operates sequentially over time, without the need for an acausal iterative settling process to produce the model output.

### Arguments for temporal prediction in the cochlea

Evolution by natural selection maximizes reproductive success (Grafen, 2014; Levin and Grafen, 2019). However, individual physiological systems of an organism can be seen as being approximately optimized for some more specialized objective or set of objectives (subject, of course, to certain constraints) that serves this ultimate objective. There are several arguments as to why the functional properties of sensory structures, such as the cochlea, may be governed by temporal prediction. First, efficient temporal prediction may induce encoding of ‘underlying’ features behind the stimuli (Bialek et al., 2001) – the requirement to accurately predict future input should help to establish an accurate neural model of the external world. Second, a sensory representation optimized for temporal prediction will disregard information in the input stimulus that is not predictive of the future, and therefore arguably superfluous. This can provide an initial filter to restrict the amount of information the brain has to deal with, since gathering and transmission of unnecessary information is energetically costly (Marzen and DeDeo, 2017). Third, sensory processing guides action, rendering the future sensory environment highly relevant; optimization for temporal prediction should enable more accurate and faster future actions, which can be critically important in a survival context. Finally, an empirical argument for temporal prediction comes from its capacity to account for the temporal and spectral characteristics of neuronal receptive fields found in diverse sensory regions and modalities, including the retina and the visual and auditory cortex (Palmer et al., 2015; Singer et al., 2018, 2019), in addition to the cochlea.

### Limitations and future development of the model

The temporal prediction model captures some important physiological features of the cochlea, but has certain limitations. The rise time of the model impulse responses is somewhat faster than that measured for the auditory nerve, and some of those responses show an initial click that is not seen in the experimental data. Also, the cochlea exhibits nonlinear properties that our model, by its design, could not capture. One example is two-tone suppression, where the simultaneous presentation of two tones with nearby frequencies nonlinearly attenuates basilar membrane responses (Robles and Ruggero, 2001). Potential developments of the model that could help capture these properties include increasing its complexity and nonlinearity to more fully enable reflection of the biophysical properties of the cochlea. In particular, adopting a recurrent network model, rather than the purely feedforward version used here, should better reflect the mechanical interactions that take place along the basilar membrane and with the outer hair cells and surrounding cochlear fluids. Furthermore, incorporating an additional layer to potentially capture the integration and adaptation at inner hair cells, auditory nerve fibers and the intervening ribbon synapses (Eatock 2000; Zilany et al., 2009; Moser and Beutner, 2000; Raman et al., 1994; Wen et al., 2012) would represent another important extension, as would introducing longer input windows and higher sampling rates in order to extend the frequency range that can be explored.

## Methods

### Auditory dataset and preprocessing

We used a dataset composed of natural sounds: birdsong and other animal calls, environmental sounds, and human speech by children and adults as well as baby vocalizations. Sound recordings of birdsong and mammal and insect calls were drawn from the Macauley Library at the Cornell Lab of Ornithology^1^ (https://www.macaulaylibrary.org/), the British Library Sounds Environment & Nature Collection (https://sounds.bl.uk/Environment) and https://animal-sounds.org/farm-animal-sounds.html. We also used a corpus of ferret vocalizations recorded in our laboratory (by Kerry Walker, Oxford Auditory Neuroscience Group) and an additional recording from https://freesound.org/people/J.Zazvurek/sounds/155115/. Recordings of adult human speech were taken from http://databases.forensic-voice-comparison.net/ (Morrison et al., 2012). We also used some human speech and environmental sounds, such as snapping twigs, which were recorded in our laboratory in an anechoic chamber. Recordings of children speaking were from the CHILDES database (MacWhinney, 2000; Edwards and Beckman, 2008) from TalkBank (talkbank.org). Baby vocalizations were from the OxVoc database (Parsons et al., 2014).

Each recording was resampled to 22,050 Hz and convolved with a filter that mimics the transformation of raw sound by the middle ear. This filter was provided by the Python package “Brian Hears” (Fontaine et al., 2011). To lessen edge effects, we also applied high- and low-pass filters (5^th^-order Butterworth) with cut-off frequencies of 500 Hz and 5,512.5 Hz, respectively. The high-pass filter was applied because the time window of the network input limited the capacity to capture frequencies with a period greater than this time window. The high-pass filter provided a smooth fall-off in power near this cut-off to avoid potential edge effects. The low-pass filter was applied to provide a smooth fall-off near the Nyquist frequency to avoid potential edge effects and because auditory nerve fibers do not phase lock at high sound frequencies.

We then divided each of the sound clips into consecutive snippets of 6.53ms (144 samples) duration. We excluded sound snippets with a root mean square value below a threshold (0.105) to avoid silent sound clips as training input. Each individual sound snippet was then normalized have a standard deviation of one. Finally, we added Gaussian noise to the input of the sound snippets at an SNR of 9 dB typically, but different values were also explored. For training and testing the temporal prediction and autoencoder models, the selected snippets were divided into a training set of *N* = 2,038,187 snippets, and a test set of 164,193 snippets.

For Supplemental Fig. 2G-H, we used a larger dataset where we selected snippets from the same clips as before, but the snippets overlapped, with the starting points of the 6.53 ms long snippets spaced by 0.65 ms. Otherwise pre-processing of the larger dataset was exactly the same as for the standard dataset.

### The cochlear temporal prediction model

The cochlear temporal prediction model was trained to predict the future of sound snippets from their past. The model consisted of a feedforward artificial neural network with one hidden layer with a nonlinear activation function. It had *i* = 1 to *I* = 128 input units, one for each time step of the input, *j* = 1 to *J* = 64 hidden units, and *k* = 1 to *K* = 16 output units, one for each time step of the target.

The *N* = 2,038,187 sound snippets of the training set were used to train the model. Each snippet, *n*, was *I*+*K* = 144 time steps (samples) long. The first *I* = 128 time steps (5.80 ms) of a snippet were taken as the past and provided the input unit activity *u*_*in*_ to the model, indexed from *i* = 1 to *I*. The last *K* = 16 time steps (0.73 ms) of a snippet were taken as the future and provided the target values *y*_*kn*_ that the model aimed to predict, indexed from *k* = 1 to *K*.

The activation *x*_*jn*_ of each hidden unit of the model was a linear mapping from the input unit activity, and was given by,

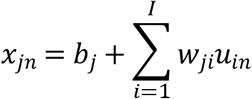

where *b*_*j*_ is the bias value for hidden unit *j, w*_*ji*_ is the weight from input unit *i* to hidden unit *j*, and *u*_*in*_ is the activity of input unit *i* for snippet *n*.

The activity *a*_*jn*_ of hidden unit *j* for training example *n* was then given by applying an activation function to the hidden unit’s activation,

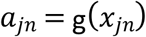

where the scaled hyperbolic tangent *g*(*x*) = tanh (*αx*)*β* was used as the activation function of the hidden units. Scaling factors *α* and *β* were set to the standard values of 2/3 and 1.7159, respectively (LeCun et al., 1998).

The model’s prediction, *v*_*kn*_, of the target, *y*_*kn*_ (the future), was provided by a linear mapping from the hidden unit activity, and was given by

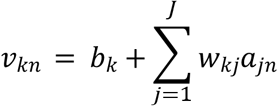

where *w*_*kj*_ is the weight from hidden unit *j* to output unit *k*, and *b*_*k*_ is the bias for output unit *k*.

The trainable parameters, *w*_*ji*_, *b*_*j*_, *w*_*kj*_ and *b*_*k*_, were optimized to perform temporal prediction by minimizing the objective function

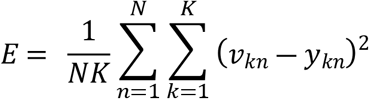

where *N* is the total amount of training snippets, and *y*_*kn*_ is the target value for output unit *k* for training example *n*. The optimization was performed using stochastic gradient descent on mini-batches of 100 snippets and the update function Adaptive Moment Estimation (Kingma and Ba, 2015).

### The cochlear autoencoder model

The cochlear autoencoder model was exactly the same as the cochlear temporal prediction model, except that it has *K* = *I* = 128 output units and that it aimed to estimate its own input. Instead of the target being y_*kn*_, the future of a snippet, it was *u*_*kn*_, the past of a snippet, which is the same as *u*_*in*_, the input to the model.

### Impulse response analysis

Hidden unit impulse responses with a power value of less than 1% of the mean impulse response power for all hidden units in the models were removed from analysis. The impulse response characteristics were found by determining the spectrum using the Fast Fourier Transform. The center frequency was defined as the frequency bin containing the spectral peak for each impulse response. The bandwidth was defined as the frequency span within a 10 dB drop on either side of the center frequency of the impulse response. Only units with a drop on both sides of the spectral peak were included in the impulse response bandwidth analysis. The temporal envelope duration of each impulse response was interpreted as being the shortest temporal span that covers 95% of the total impulse response power.

The frequency glide slopes were measured by the zero-crossing method of Carney et al. (1999), using methods very similar to theirs (see Supplemental Figure 1). First, the impulse response was smoothed. To do this, the first sample of the impulse response was excluded (due to its click characteristics) and the impulse response was padded with 5.8 ms of zeros on either side. This was then filtered forwards and backward in time with a 4^th^ order Butterworth bandpass filter, and the section corresponding to the impulse response extracted. The net result of this process produced the impulse response filtered by an 8^th^ order bandpass filter with zero phase, and hence with no introduced time delay. The bandwidth of the 4^th^ order Butterworth filter was one octave, centered at the unit’s center frequency. The envelope of the filtered impulse response was measured by taking its Hilbert transform and a 3-point average, and the time range for which the amplitude of the envelope was within 12 dB of the peak was found. Next, the mean (DC) value was calculated for a span that was the time range rounded down to an integer number of cycles of the centre frequency, centered at the midpoint of the range. This DC value was subtracted from the impulse response. The impulse response was then linearly interpolated by a factor of 1000, any zero crossings within the time range were found, and the time difference between consecutive zero crossings determined. The reciprocal of this time difference is the instantaneous frequency at the midpoint of the two timepoints. The instantaneous frequency was plotted as a function of these midpoints, a straight line was fitted to this plot by least squares, and its slope was measured. This was done for each impulse response that included at least two periods within the 5.8 ms span of the impulse response and which had a center frequency of less than 4.6 kHz, since this was the highest characteristic frequency of the auditory nerve fibers recorded by Carney et al. (1999). An additional reason for limiting this frequency range is that the zero-crossing method of measuring instantaneous frequency is inaccurate above half the Nyquist frequency, 5.5 kHz (Hoeks et al., 1984).

## Computational implementation

Custom code was written in Python for data handling, modelling and data analysis, and run on NVIDIA GeForce GTX-1080 GPUs.

## Acknowledgements

We thank Laurel Carney for the data in Figure 3B. We are grateful for the funding provided by a Clarendon Fund Graduate Scholarship to FT and a Wellcome Principal Fellowship to AJK (WT108369/Z/2015/Z).

---

**Supplemental Figure 1.**
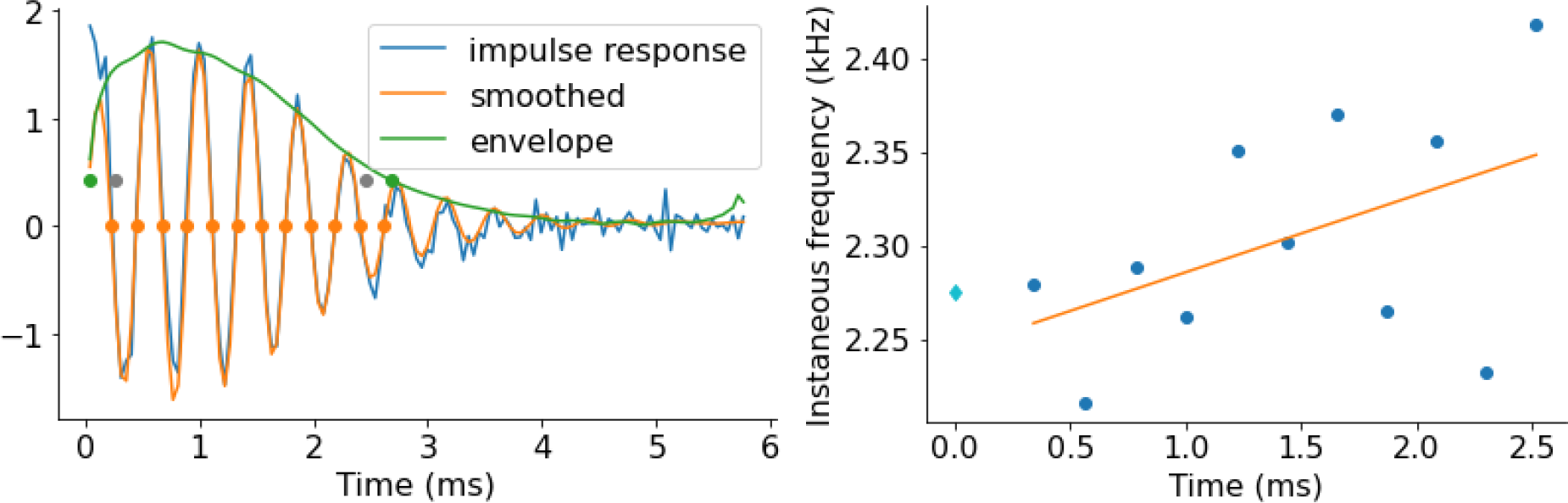
Measuring the slope of the instantaneous frequency for an impulse response. **(A)** Plot of the impulse response (blue), the smoothed impulse response (orange) after filtering and DC removal, and the envelope (green). The green dots are the edges of the range over which the zero crossings were measured, where the envelope amplitude is 12 dB down from the envelope peak. The gray dots indicate the range over which the DC was measured, an integer number of periods of the center frequency between the green dots. The orange dots are the zero crossings. **(B)** Blue dots show the instantaneous frequency against time of the impulse response in A. The instantaneous frequency is the reciprocal of the time difference between consecutive zero crossings. The cyan diamond shows the center frequency of the impulse response. The orange line is the best linear fit, which provides the slope. The slopes of each unit were then used to make Figure 4F.

**Supplemental Figure 2.**
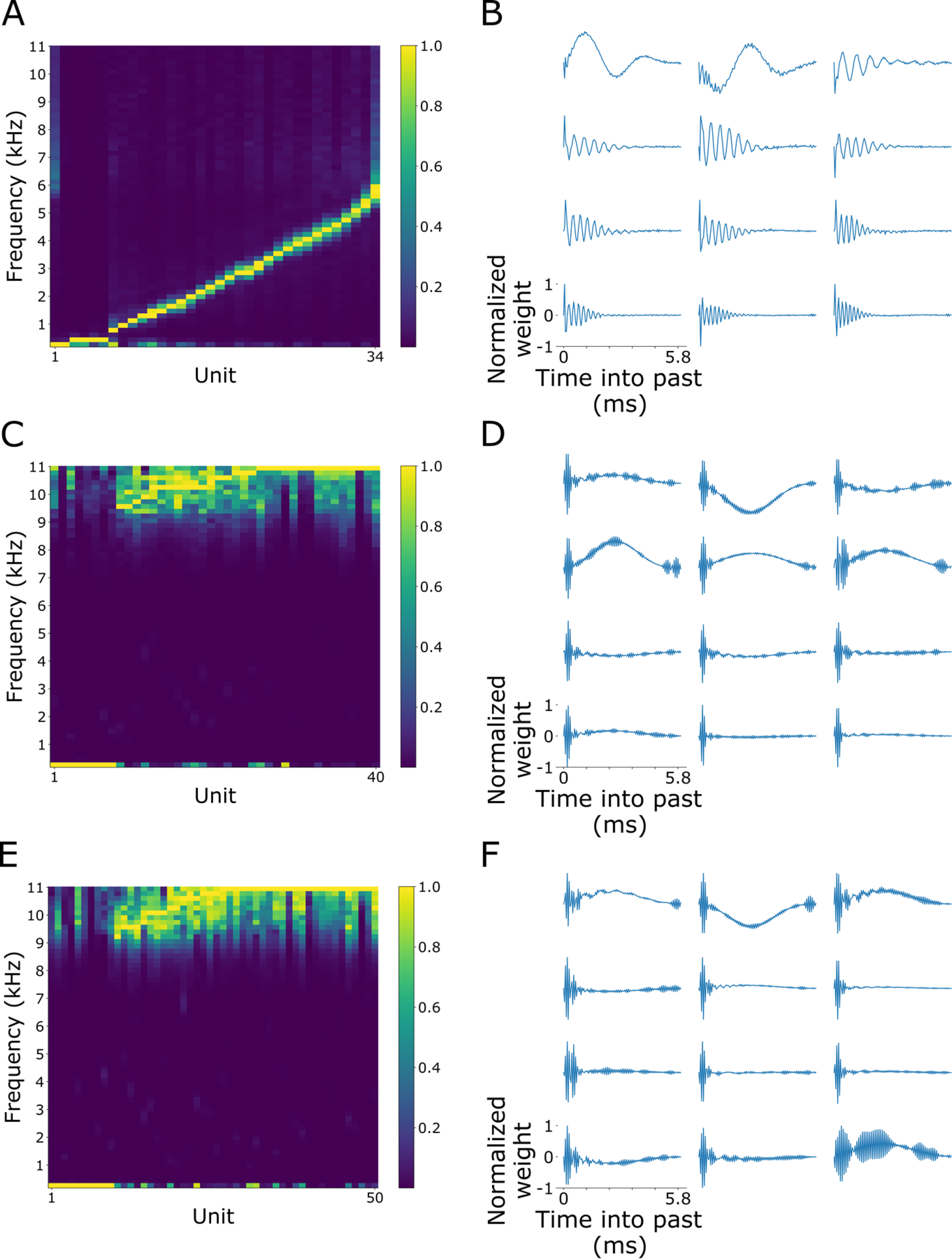
Impulse response characteristics of the temporal prediction model trained on different noise schemes. **(A, B)** Noisy input and target. **(C, D)** Noisy target. **(E, F)** Large noiseless dataset. **(A, C, E)** The normalized power spectrum of the impulse response of the hidden units of the feedforward model. The color bar represents the normalized power. If the power of the impulse response failed to exceed a threshold of 1% of the average model impulse response power, it was removed from analysis (30, 24 and 14 hidden units, respectively). **(B, D, F)** Impulse responses of the network model. The impulse response for each hidden unit was normalized to its respective maximum absolute value.

**Supplemental Figure 3.**
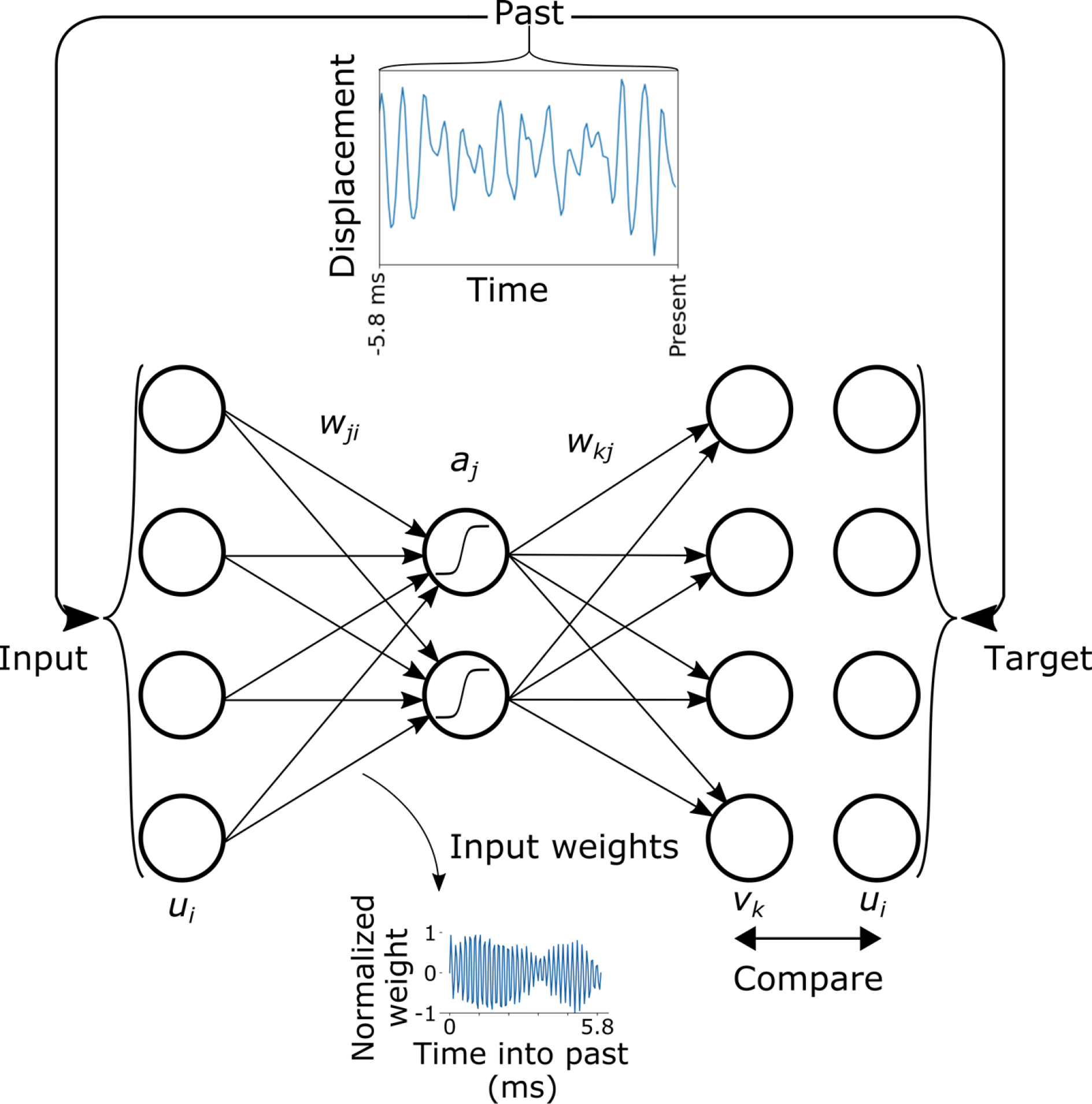
Schematic of the autoencoder model of the cochlea. The same dataset of sound snippets used to train the temporal prediction model was provided as inputs to a feedforward autoencoder neural network, and the network was optimized to recreate its input. More specifically, the weights, *w*_*ji*_ and *w*_*kj*_, and the biases, *b*_*j*_ and *b*_*k*_, of the autoencoder were optimized to recreate the input *u*_*i*_ using estimate *v*_*k*_, for the large dataset of natural sounds. The upper inset figure shows an example sound snippet representing the past. This 5.8 ms duration snippet, *u*_*i*_, served as both input and target for the autoencoder objective. The activity of an input unit represents one sample of the 128 samples constituting the snippet, while each output unit represents a particular sample of the model’s recreation of the same 5.8 ms of the stimulus. An example of the tuning properties learned by the model is shown in the lower inset figure. The incoming weights, *w*_*ji*_, to hidden unit *j* can be interpreted as the impulse response of this hidden unit.

**Supplemental Figure 4.**
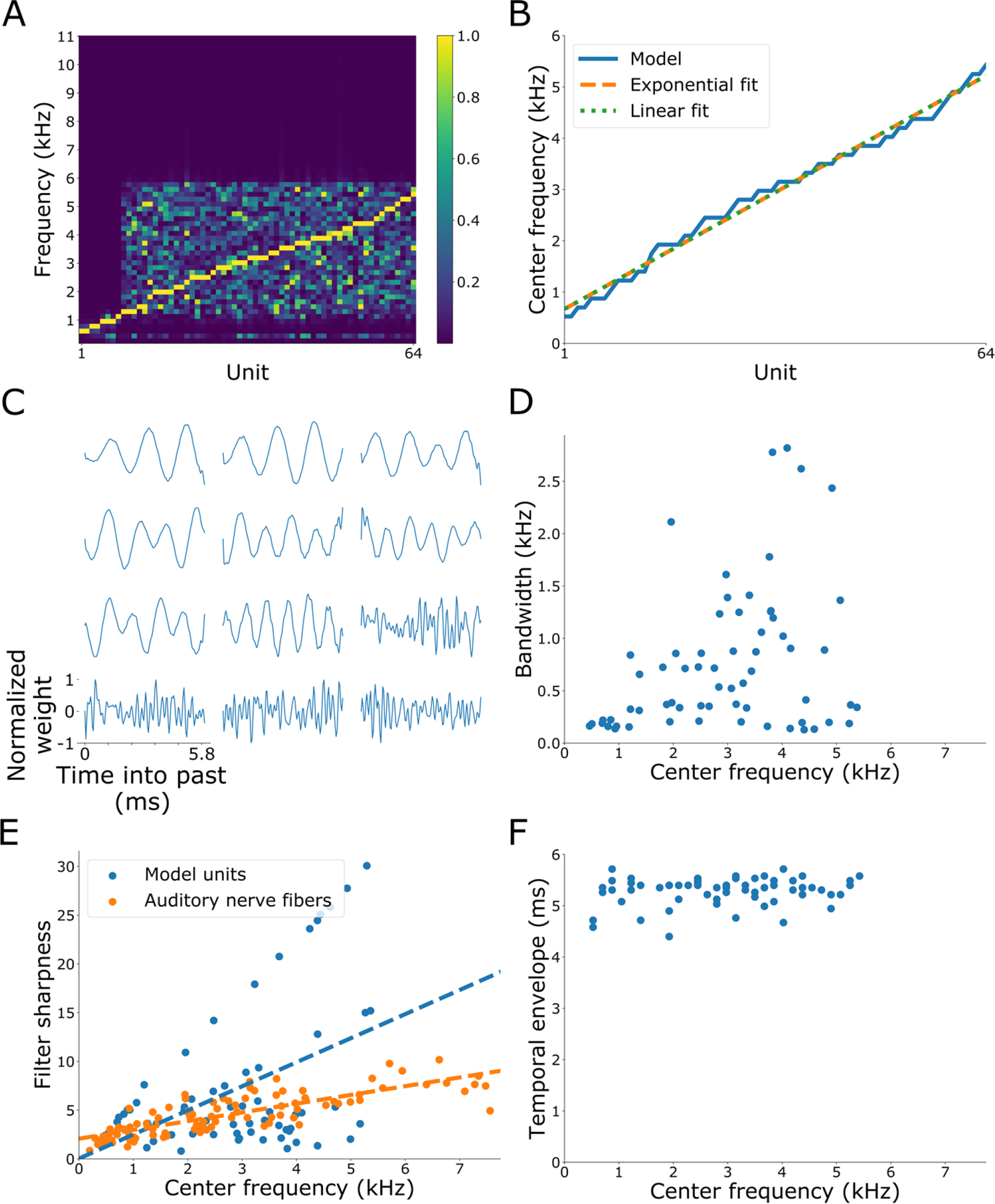
Autoencoder model trained on noiseless sound. **(A)** The normalized power spectrum of the impulse response of the hidden units of the feedforward model. The color bar represents the normalized power. **(B)** The center frequency of each model unit as a function of hidden unit rank. The hidden units are interpreted to span a contiguous section of the basilar membrane in tonotopic order, with each unit evenly spaced along this section. The model center frequencies were fitted with a logarithmic equation that has been used to describe the relationship between distance along the basilar membrane and characteristic frequency (Greenwood, 1961, 1990, 1996; Robles and Ruggero, 2001). The best linear fit of the model centre frequencies is also shown. The linear equation fitted the centre frequencies of the model units better than the exponential equation (F test, *F*(62,61) = -0.0159, *p* = 1). **(C)** Impulse responses of the network model. The impulse response for each hidden unit was normalized to its respective maximum absolute value. **(D)** The impulse response bandwidth of each hidden unit plotted against centre frequency (Pearson’s correlation coefficient, ρ = 0.326, *p* = 0.009). The impulse response bandwidth was estimated as a 10 dB decrease on either side of the spectral peak of the impulse response. **(E)** The filter sharpness of each hidden unit plotted against center frequency (ρ = 0.45, *p* = 0.0002). The filter sharpness was determined by dividing center frequency by the impulse response bandwidth. Blue dots indicate data points taken from the temporal prediction model. Orange dots indicate data points taken from cat auditory nerve fibres (ρ = 0.816, *p* = 0.0002) (Evans, 1975). **(F)** The temporal envelope of each hidden unit plotted against center frequency (ρ = 0.182, *p* = 0.15). The temporal envelope of each impulse response was determined by the shortest time window possible that captured 95% of the power of the impulse response.

## Footnotes

1 Specifically, we use the following recordings from the Macauley Library: ML3429, ML3891, ML53181, ML55348, ML55350, ML59291, ML61494, ML62970, ML67895, ML70539, ML72995, ML80731, ML100723, ML100797, ML100857, ML111595, ML116302, ML116303, ML126289, ML129248, ML130909, ML132511, ML132526, ML136504, Ml164696, ML171372, ML191165, ML191178, ML191290, ML197064, ML199078, ML201215, ML205481, ML205774, ML207517, ML206448, ML207181, ML210645, ML213055, ML213356, ML217313, ML217490, ML220049, ML233387, ML516692, ML527184 and ML527292.

## Notes

### Competing Interest Statement

The authors have declared no competing interest.

